# Accurate determination of meat mass fractions using DNA measurements for quantifying meat adulteration

**DOI:** 10.1101/2020.06.14.150375

**Authors:** Sasithon Temisak, Pattanapong Thangsunan, Jiranun Boonil, Watiporn Yenchum, Kanjana Hongthong, Teerapong Yata, Leonardo Rios-Solis, Phattaraporn Morris

## Abstract

The problem in meat adulteration and food fraud emphasised the requirement of developing accurate analytical approaches for the quantitative detection in helping the control of meat adulteration. In this study, the droplet digital Polymerase Chain Reaction (ddPCR) assays to quantify the ratios of pork DNA to the total amount of meat DNA were developed by challenging against DNA extracted from a range of gravimetrically prepared matrices of pork in beef. A single copy nuclear DNA gene, *β-actin*, was employed as a target gene, accompanied with *myostatin* gene as a cross species target for mammal and poultry meat background in order to quantifying approach. All the developed assays, singleplex, duplex and triplex did not show significant difference in quantification of pork content in beef background and demonstrated a good and comparable performance to the mass fractions. The singleplex assay provided more biases than the other two assays when performing with a low concentration of target species. The duplex assay provided a simultaneous quantification of pork and *myostatin*, whereas the triplex assay was able to detect pork, beef and *myostatin* with a decrease of technical error, cost and running time. All proposed methods allowed us to quantify pork addition in beef with a limit of quantification (LOQ) estimated at 0.1% (w/w) and a limit of detection (LOD) down to 0.01% (w/w). The developed triplex assay was also tested with commercial processed foods and showed the ability to determine not only the presence of particular pork or beef but also the quantitative purpose directly without standard curves. Hence, the developed ddPCR assays demonstrated a good trueness and precision of the methods in quantifying pork or beef content for meat adulteration. It is expected that these developed approaches can be applied to help regulators to confidently enforce food labelling obligations.

## Introduction

Meat adulteration and food fraud have recently become a globally widespread problem. According to the US Federal Meat Inspection Act (FMIA) and the European Parliament under Regulation (EC) No. 178/2002, adulteration of meat products with undeclared meat species is not allowed (Grundy et al. 2013). This includes the substitution of higher-valued meats with lower-valued ones during production process, which can lead to an unfair competition (Fajardo et al. 2008). The presence of undeclared species may cause health problems such as food allergy (Naaum et al. 2018; Temisak et al. 2019) and a fatal neurodegenerative disease caused by bovine spongiform encephalopathy (Sultan et al. 2004). The previous studies reviewed that over 50% of processed meat products tested were mislabeled (Cawthorn et al. 2013; Di Pinto et al. 2015; Quinto et al. 2016). Therefore, accurate methods for determination of meat species in food products are required.

There are several analytical methods used for identifying meat species and most of which are based on protein and DNA detection. Protein-based methods such as electrophoretic techniques (Montowska and Pospiech 2007), chromatography-mass spectrometry (Grundy et al. 2008) and enzyme-linked immunosorbent assay (ELISA) (Chen and Hsieh 2000; Hsieh and Ofori 2014) have been successfully employed for detecting meat species. These techniques, however, showed some disadvantages such as imprecision as well as less specificity and sensitivity as proteins can be denatured by heat, pressure, exposure to high heavy metal and salt concentrations. Therefore, the methods that rely on protein measurements may not be suitable for processed-food products and also for quantitative purpose. On the other hand, DNA-based methods are more stable, easy to operate, and widely used with a broad range of detections (Martin et al. 2009). Quantitative real-time Polymerase Chain Reaction (qPCR) is the technique based on DNA detection and has been used for detecting a range of meat species (Köppel et al 2020; Soares et al. 2010). This technique is highly sensitive, sequence-specific, amendable to a type of samples and has a vast dynamic range of detection (Nixon et al. 2015). However, the need for reference standards to generate a calibration curve is one of the downsides of qPCR (Ren et al. 2017). Recently, digital PCR (dPCR) has been adopted for quantifying the amount of nucleic acid targets by partitioning them into a number of reaction chambers (Whale et al. 2016). As a result, each reaction contains one or no copy of nucleotide sequences of interest that can be assayed individually and counted without requiring a standard curve and correcting by Poisson statistics (Baker 2012). Compared to qPCR, dPCR is more advantageous in terms of its sensitivity, specificity and precision (Baker 2012; Köppel et al 2019; Manoj 2016; Ren et al. 2017; Cao et al. 2020, Wang et al. 2018).

The main focus of this work was not only on the qualitative detection but also the quantification. Many previous studies have the problem in quantification of meat by converting the DNA measuring to the weight proportion of meat (Cai et al. 2017; Martin et al. 2009; Soares et al. 2010). The known type of meat is the key factor in quantifying the meat content by the DNA measuring method. To calculate the percentage of the target meat in meat background, several assays and calibration curves may be needed to quantify the background meats as the background meats can be pork, beef, goat, lamb, chicken or turkey. Moreover, the DNA measurements may not reflect the actual quantity (mass fractions) if the selected target are multiple genes such as mitochondrial gene (Nixon et al. 2015). Ren and colleagues tried to use the *k* constant to transform the copy number ratio to the mass fraction by ddPCR method (Ren et al. 2017). However, the known types of meat were also required in order to use the correctly *k* constant and assays to determine meat species in the fact that meat products in markets may contain with an unknown and variety of meat type, resulting in missed quantification.

In order to tackle the previously mentioned challenges, in this study, singleplex, duplex and triplex ddPCR assays for accurate mass determination of pork present in a sample of mixed pork and beef matrices were developed and validated. To achieve the accurate quantification of pork mass percentage in meat background, the selected single copy nuclear DNA target for species identification together with a selected single copy cross species gene for a broad meat species background (mammal and poultry) was used for ddPCR. We found that pork or beef DNA fractions were consistent with the gravimetric proportions (w/w) with an acceptable level of precision and trueness.

## Materials and Methods

### Sample Preparation

Fresh pork (*Sus scrofa*), processed foods, and non-target animal species including chicken (*Gallus gallus*), duck (*Anas platyrhynchos*), partridge (*Perdix perdix*), salmon *(Salmo salar)*, and crocodile (*Crocodylus siamensis*) were purchased from supermarkets in Pathum Thani, Thailand while beef (*Bos taurus*) were obtained from halal food markets in Pathum Thani, Thailand. In the control groups, raw pork and beef were separately blended, freeze-dried (CHRIST, Gamma 1-16 LSC, Germany), sieved through of 300-µm standard test sieves (Retsch, Fisher scientific, USA), and stored at −80 °C until further use. To mimic highly processed foods, raw pork and beef were separately autoclaved for 20 min at 121 °C and then processed with the same methods for raw meat. For raw meat matrix mixtures, pork meat powders were prepared at the 100%, 75%, 50%, 25%, 10%, 1%, 0.5%, 0.1%, 0.01% and 0% (w/w) levels in beef and for autoclave treated meat, 100%, 1% and 0% pork in beef (w/w) were prepared by gravimetric balance (Mettler Toledo, XPE26, Switzerland). Eleven independent replicates were prepared for the limit of detection (LOD) and the limit of quantification (LOQ) determination.

### DNA extraction

Genomic DNA was extracted and purified from 2 g of samples with the cetyl trimethylammonium bromide (CTAB) method and Genomic-tip 100/G (Qiagen, Hilden, Germany) according to the standard DNA extraction method (EU-RL 2013). The quality and concentration of each extracted DNA were measured using a Microplate Reader (Tecan Spark, Switzerland). The quality of extracted DNA were assessed using 1.5% agarose gel electrophoresis.

### Oligonucleotide primers and probes

Primers and probes used in this work have been published elsewhere (Köppel et al. 2011; Laube et al. 2003). All the primers and probes were synthesised and purified by Macrogen Inc. (Seoul, Korea). The nucleotide sequences of the primers and probes, and PCR product sizes are presented in Table 1.

**Table 1.**
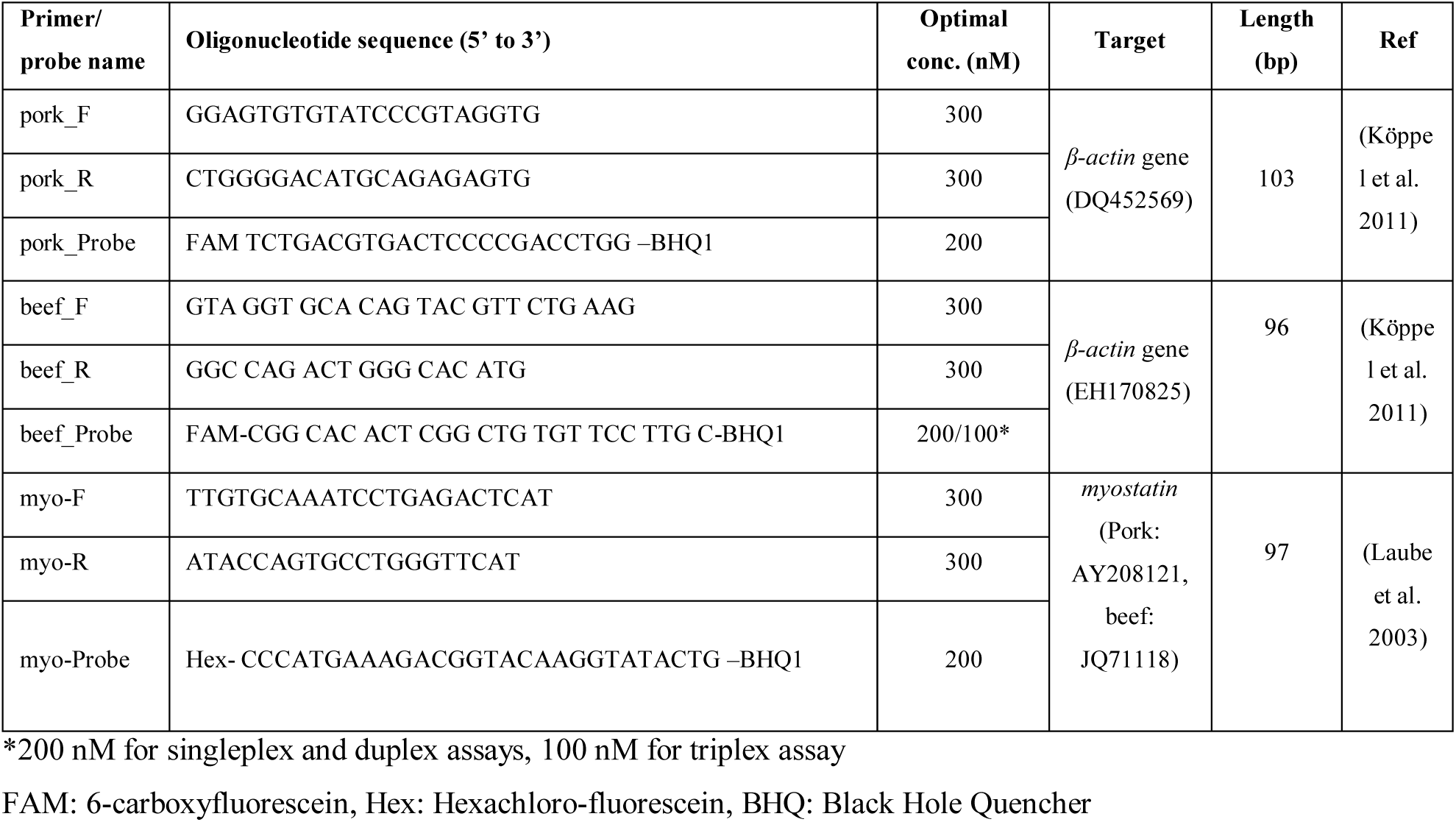
Primers and probes used in this study

### Quantitative real-time PCR

The qPCR reactions were performed in 20 μL comprising 1x TaqMan Universal PCR mastermix (Applied Biosystems, USA), optimal concentrations of forward and reverse primers, 200 nM of probe, approximately 30 ng of DNA template and water. The thermo-cycling of qPCR (ABI 7500 Real-Time PCR) was performed at 95 °C, 10 min for enzyme activation, followed by 45 cycles of 15 sec at 95 °C of denaturation and 1 min at 58 °C (optimal temperature) for annealing and extension. qPCR data was analysed by 7500 Software v2.3 (ABI 7500 Real-Time PCR). Primer and probe concentrations were optimised according to suggestion in the manufacturer’s instructions (TaqMan Universal PCR mastermix, Applied Biosystems, USA).

### Droplet digital PCR

To perform ddPCR, the prepared Mastermix for ddPCR contained 1XddPCR supermix probes without dUTP (Bio-Rad, USA), 300 nM (optimal concentrations) of each primer, 200 nM of probe, 5U of *Hind*III, DNA template (approximately 1,000 copies/µL of sample) and water to make up 20 µL of its total volume. The combinations of the assays were Pork/Myo and Pork/Beef/Myo assays for duplex and triplex experiments, respectively. Only in triplex reaction, beef probe concentration was reduced to 100 nM. The PCR reactions were prepared in a 96-well plate (Bio-Rad, CA, USA). The plate was placed on a AutoDG (Bio-Rad, USA) for droplet generation. After generation of the droplets, the collected droplets plate was sealed using pierceable foil heat seal and the PX1 PCR plate sealer (Bio-Rad, CA, USA) at 180 °C for 5 sec. The PCR cycle of ddPCR was performed at 95 °C for 10 min for enzyme activation, 40 cycles of 30 sec at 94 °C for denaturation, 1 min of 58 °C (optimal temperature) for annealing and extension, and one cycle at 98 °C, 10 min for enzyme deactivation in a thermal cycler (T100, Bio-Rad, Pleasanton, CA, USA) with the temperature ramp rate of 2 °C/s. To obtain the optimal annealing temperature for ddPCR assays, the gradient temperature between 55-64 °C was used. The optimal annealing temperature of 58 °C was chosen. When the PCR was completed, the droplets were read using a QX200 Droplet Reader (Bio-Rad, CA, USA). The data was finally analysed by QuantaSoft software (v1.7.4.0917, Bio-Rad, CA, USA). The Digital MIQE guidelines were followed (Table S1).

## Results and discussion

### Method validation

In this study, *β-actin* gene was selected as a species-specific target gene for pork and beef identification (Köppel et al. 2011). *β-actin* gene is found as a single copy in pork and beef genomes. For quantitative measurements, therefore, the use of *β-actin* gene is more suitable than multiple copy number genes such as mitochondrial DNA and 18s rRNA genes, as these genes can vary among tissue types and animal species (Barakat et al. 2014). In order to transform the ratio of DNA copy numbers to the mass proportion, a constant number factor has been applied (Ren et al. 2017). However, when this factor was applied to the mixture of the unknown meat, the method has to be monitored for accuracy since the constant factor depends on the type of meats (Ren et al. 2017). Here, a broad range of reference genes were introduced to quantify a variety of meat DNA backgrounds. *Myostatin* gene was selected as a cross species target for measuring the total amount of meat DNA content. This was due to the single copy number and highly conserved degree of *myostatin* gene sequences amongst animal species including mammals and poultry that cover most types of meat tissues (Laube et al. 2003; Nixon et al. 2015). This gene has been previously used as the internal control in qPCR reactions, and as quality control in DNA extraction step (Laube et al. 2003).

The nucleotide sequences for primers and probes used for ddPCR in this study (Table 1) were obtained from previous works using qPCR (Köppel et al. 2011; Laube et al. 2003). The assays have been already tested for the specificity by using qPCR (Köppel et al. 2011; Laube et al. 2003; Nixon et al. 2015). In this study, the specificity of the assays were also verified by performing the qPCR with isolated DNA from pork (*Sus scrofa)*, beef (*Bos taurus*), chicken (*Gallus gallus*), duck (*Anas platyrhynchos*), partridge (*Perdix perdix*), salmon *(Salmo salar)*, crocodile (*Crocodylus siamensis*) and Human *(Homo sapiens)* (Promega, USA), while water was used as NTC. The qPCR result indicated that pork and beef assays were specific to their target species (Figure S1A and S1B). The myo assay showed positive detection with DNAs from mammals and poultry whereas the assay did not show positive detection for DNAs from salmon *(Salmo salar)* and crocodile (*Crocodylus siamensis*) (Figure S1C). This suggested that the *myostatin* is a potential cross species gene for assaying the mammal and poultry DNA.

The ddPCR method is generally more robust than qPCR method, however, the fluorescence amplitude and annealing temperature still need to be optimised for a clearly separation of the clusters between positive and negative droplets (ISO-20395 2019). In this study it was found that the optimal concentrations of the forward and reverse primers were 300 nM, while the optimal concentrations of the probes were 200 nM for all the assays. In triplex assay, the optimal probe concentration for the beef assay was 100 nM (Table 1). The optimal annealing temperature for ddPCR was approximately 58 °C (Figure S2) in which the signal obtained from positive droplets was clearly separated from negative droplets. In this study, the PCR efficiency of pork and beef assays was 94.93% ± 3.50% (Figure S3 A and E) and 96.38% ± 1.54% (Figure S3 B and F), respectively, while that of the myo assay with pork DNA was 97.23% ± 5.16% (Figure S3 C and G) and the myo assay with beef DNA was 101.58% ± 4.34% (Figure S3 D and H).

### Singlexplex, duplex and triplex assays

In the development of ddPCR assays for pork and beef DNA detection and quantification, three ddPCR assays, singleplex, duplex and triplex, were tested cross reactivity and compared. In the singleplex assay, pork, beef and myo assays were independently performed. The pork assay showed positive detection when tested with the samples containing pork DNA (Figure 1A), whereas no signal was detected with the samples containing only pure beef DNA (Figure 1B). On the other hand, the beef assay showed a positive signal for the beef DNA samples (Figure 1C), but undetectable results for the samples with pork DNA only (Figure 1D). As expected, the myo assay for *myostatin* gene showed a positive detection for pork and beef DNA samples (Figure 1E and F).

**Figure 1.**
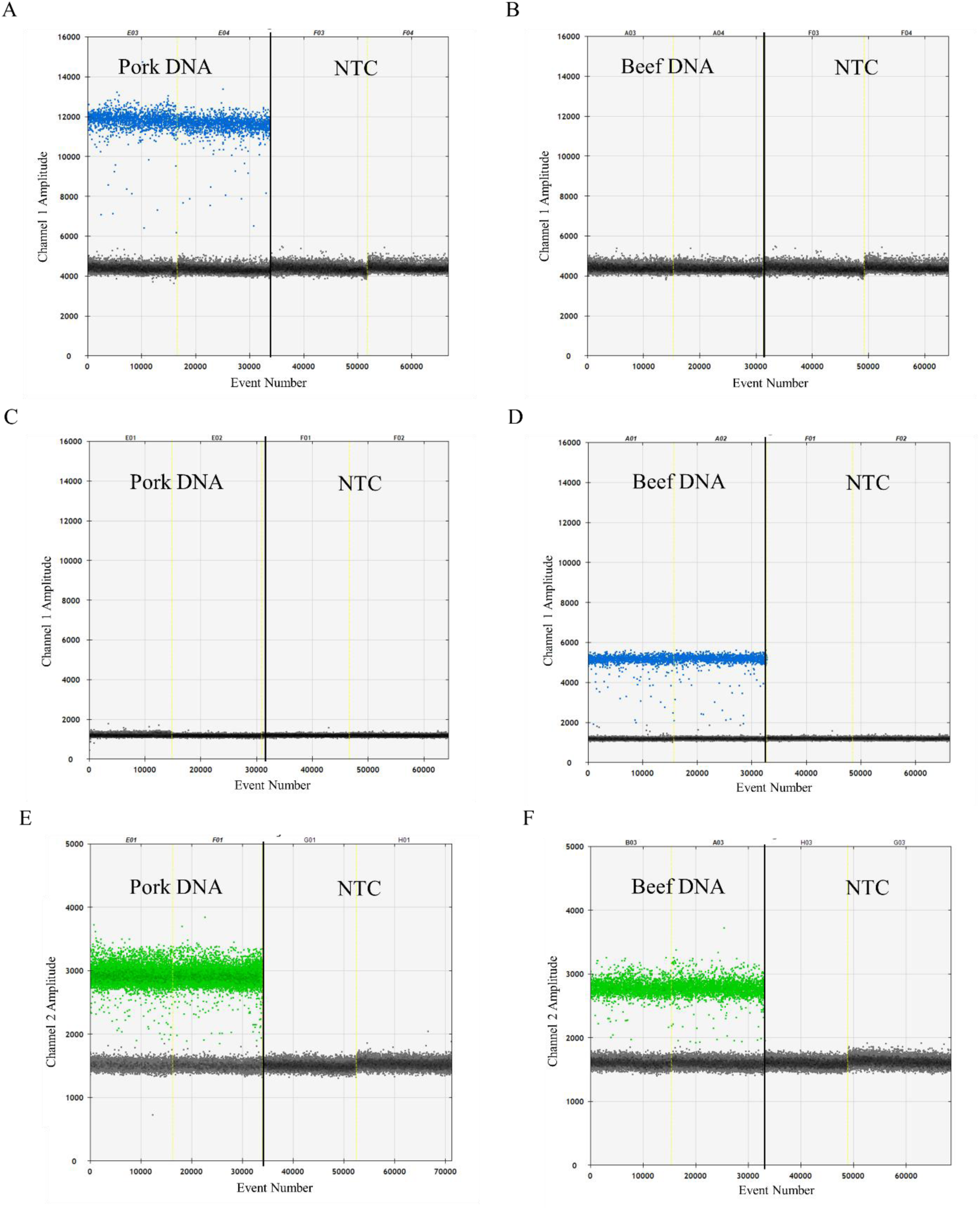
Singleplex ddPCR assays for identification of pork and beef gDNAs. The representatives for each assay were taken and presented as follows: (A) pork DNA with pork assay, (B) beef DNA with pork assay, (C) pork DNA with beef assay, (D) beef DNA with beef assay, (E) pork DNA with the myo assay and (F) beef DNA with myo assay. NTC is no-template control (ddH_2_O). Black dots indicate negative droplets, blue dots are positive droplets for beta-actin genes, and green dots are positive droplets for *myostatin* genes.

Since the singleplex assay is time-consuming and cost-ineffective (Whale et al. 2016), duplex and triplex ddPCR assays were performed for the quantification of pork and beef to improve such issues. In the duplex assay, two genes, pork or beef *β-actin* and *myostatin*, were simultaneously detected. Figure 2 shows the results of ddPCR for pork-myo and beef-myo assays. Using pork-myo duplex assay to detect the samples containing 100% purified pork DNA, the assay was able to detect pork *β-actin* and *myostatin* at the same time (Figure 2A). In contrast, when pork-myo duplex assay was performed with 100% purified beef DNA, only the *myostatin* was detected (Figure 2B). In the same way, the beef-myo duplex assay was tested for its usability. Results showed that only *myostatin* gene was detected in the 100% pork DNA samples (Figure 2C). Positive signals from both beef *β-actin* and *myostatin* can be observed when 100% beef DNA was used (Figure 2D).

**Figure 2.**
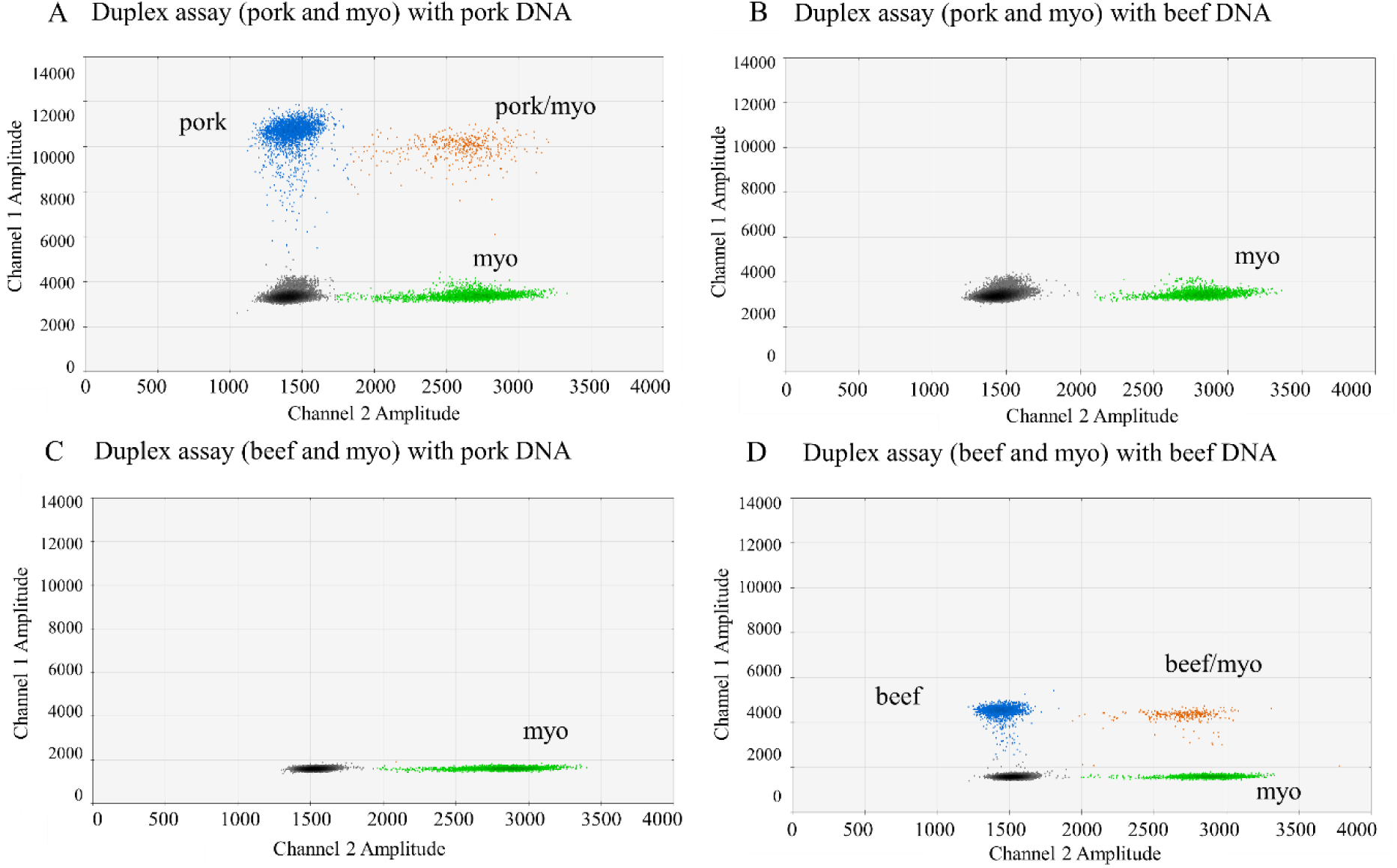
The 2D plots for duplex ddPCR assays for quantifying pork and beef genomic DNA. Each figure is (A) pork-myo duplex with 100% (w/w) pork DNA, (B) pork-myo duplex with 100% beef DNA, (C) beef-myo duplex with 100% pork DNA, and (D) beef-myo duplex with 100% beef DNA. Black dot populations are negative droplets, blue dot populations are positive droplets for pork or beef β-actin, green dot populations are positive droplets for *myostatin*, whereas orange dot populations are positive droplets for *β-actin* and *myostatin*.

In the triplex ddPCR assay, the limitation of ddPCR is that there are only two channels for two fluorescent dyes, therefore, to simultaneously detect multiple targets, varying the concentration of probes labelled with the same fluorescent dyes should be done (Whale et al. 2016). The probe concentration for the pork assay (200 nM-FAM-probe) was twice as high as the probe for the beef assay (100 nM-FMA-probe), while *myostatin* was labelled with a different fluorescence dye (Hex-labelled). The triplex assay was also tested with 100% pork DNA and showed a positive detection for pork *β-actin* and *myostatin* (Figure 3A). When performing the triplex ddPCR with 100% beef DNA, positive signals were seen for beef *β-actin* and *myostatin* (Figure 3C). To show the possible results in order to determine three independent targets test of the triplex assay, the mixture of 50% pork and 50% beef DNAs was used as a template for ddPCR. Figure 3B and D illustrate that positive droplets were found for pork *β-actin*, beef *β-actin* and *myostatin* even in one PCR reaction, indicating that the triplex assay was able to measure three target genes at a time.

**Figure 3.**
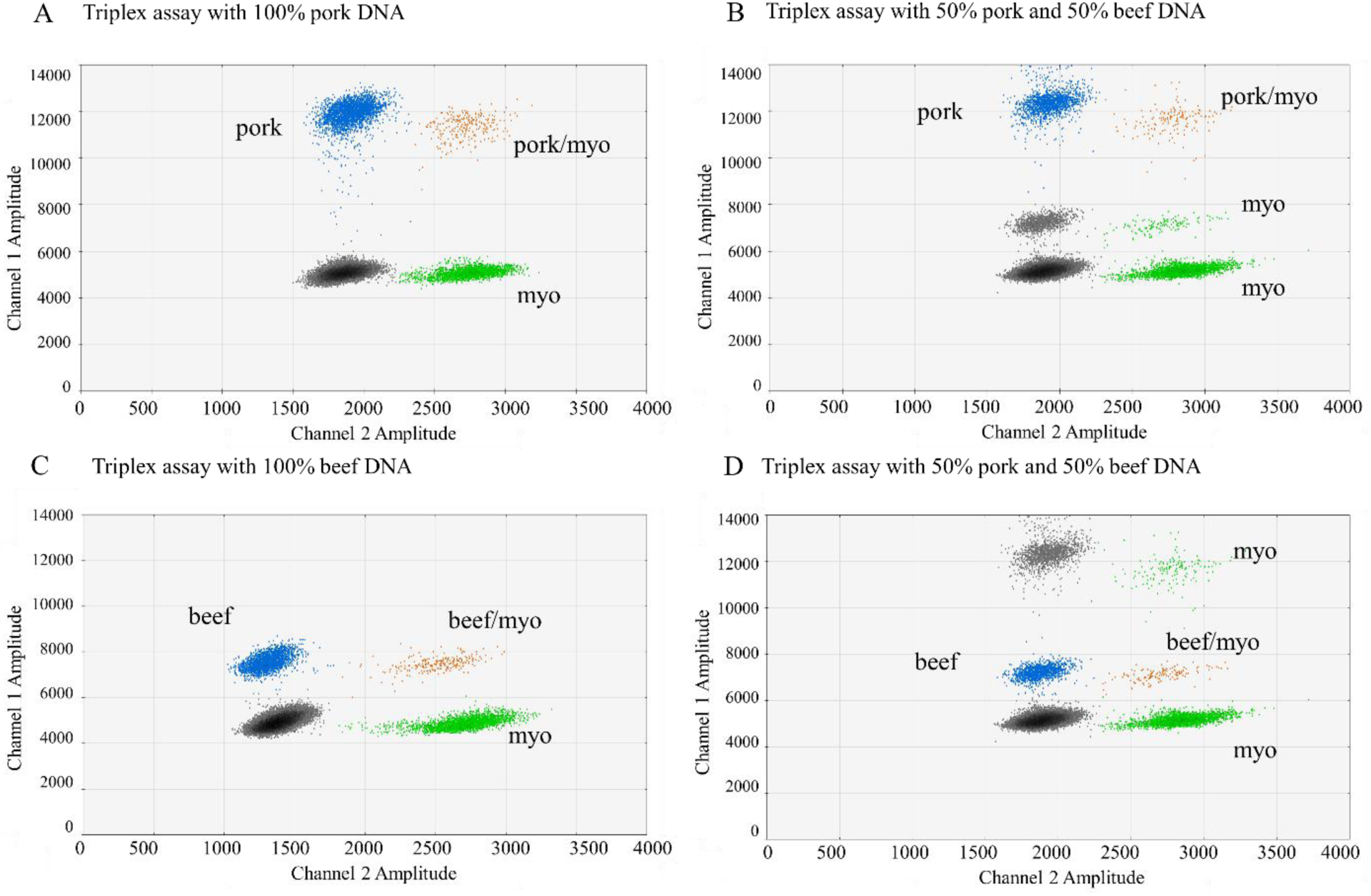
The 2D plots for triplex ddPCR assays for quantifying pork and beef genomic DNA. Each figure is triplex assays tested with (A) 100% (w/w) pork DNA, (B) 50% pork DNA and 50% beef DNA, highlighting pork *β-actin* positive detection, (C) 100% beef DNA, and (D) 50% pork DNA and 50% beef DNA, highlighting beef *β-actin* positive detection. Black dot populations are negative droplets, blue dot populations are positive droplets for pork or beef *β-actin*, green dot populations are positive droplets for *myostatin*, and orange dot populations are positive droplets for *β-actin* and *myostatin*.

### The ratio consistency of *β-actin* and *myostatin* genes

Since the key aim of our research was to investigate the potential of the selected single copy gene to quantify the mass fractions of pork in meat products, it was important to confirm the consistency of 1:1 ratio of pork or beef *β-actin* species-specific target gene to a cross species target (*myostatin*) gene. We used duplex ddPCR to confirm the consistency of the ratio between *β-actin* species-specific target and *myostatin* genes. If both genes were a single copy in the genome, the *β-actin* and *myostatin* ratio should be close to one irrespective if the concentrations of the genes changed by the serial-dilutions. The results showed that the ratio of pork *β-actin*/*myostatin* (Figure 4A) and beef *β-actin*/*myostatin* (Figure 4B) were near or equal to 1 as predicted although the concentrations of *β-actin* and *myostatin* genes were varied. Since pork *β-actin*, beef *β-actin* and *myostatin* were proved to be a single copy gene in pork or beef genome, then the pork, beef and myo assays were further used in the DNA quantification approach.

**Figure 4.**
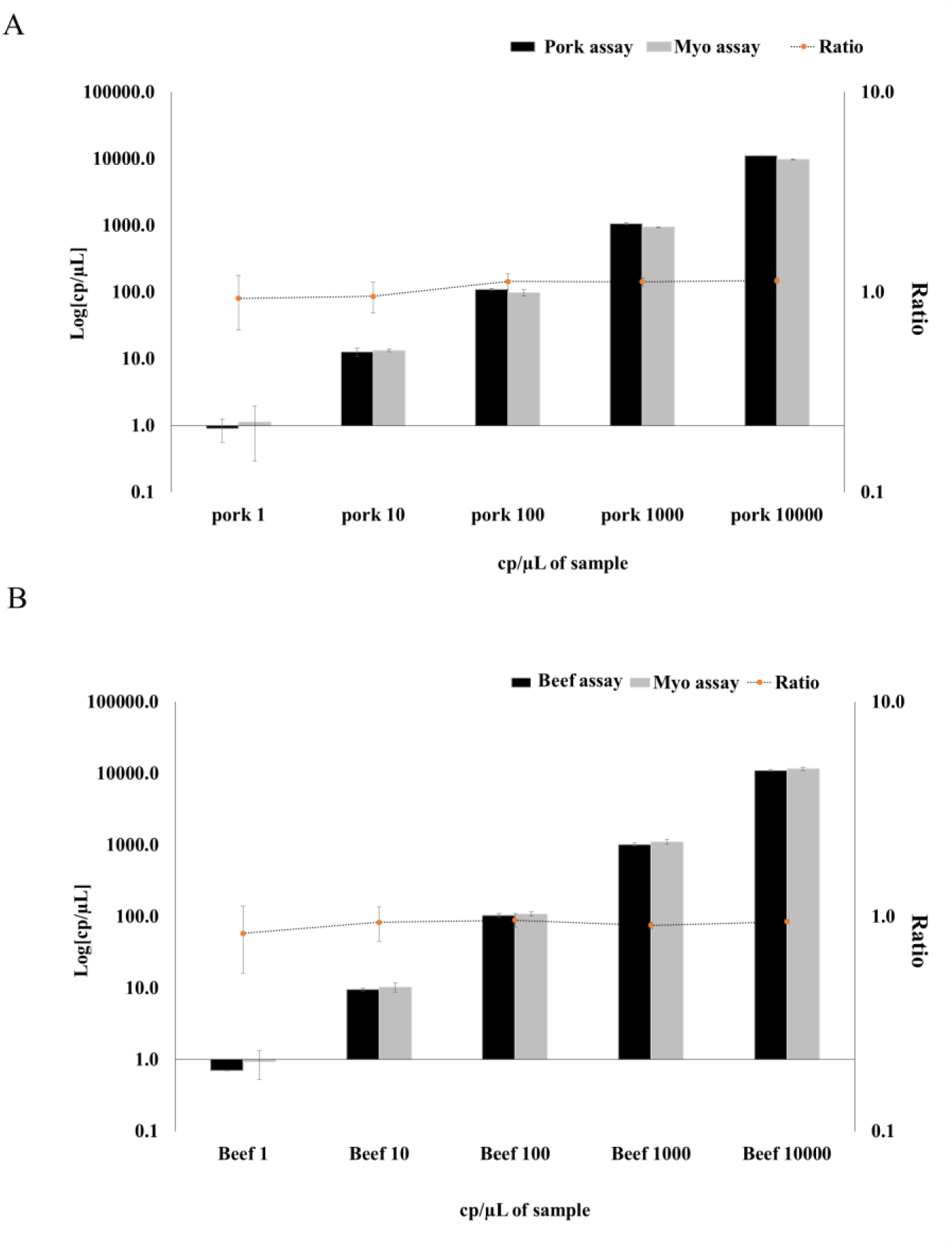
The ratio of *β-actin* and *myostatin* genes determined by duplex ddPCR assays. A tenfold dilution series (10,000 – 1 copies/µL of sample) of purified pork (A) or beef (B) DNA was assessed by duplex ddPCR (pork/myo and beef/myo assays) for absolute pork *β-actin* and *myostatin* or beef *β-actin* and *myostatin* concentrations (the left y-axis). The black bars represent the amount of *β-actin* gene as a species-specific target whereas the amount of *myostatin* gene is represented by the grey bars. The concentrations of pork or beef DNA are on the x-axis. The ratio (cp/cp) of *β-actin* gene to *myostatin* gene is represented by the dotted line with orange dots (the right y-axis). The error bars are the standard deviation obtained from three independent experiments (n=3).

### Testing the developed ddPCR assays with isolated DNA from mixed meat matrices

In this section, the copy number ratio between a species-specific target gene and a cross species gene was used to determine the mass fraction of pork in mixed pork and beef matrices. Prior to independent triplicate DNA template extractions, various ratios of mixed pork and beef matrices were gravimetrically prepared as follows: 100%, 75%, 50%, 25%, 10%, 1%, 0.5%, 0.1%, 0.01% and 0% (w/w) pork in beef. The extracted DNA samples from mixed matrices were then investigated through either singleplex, duplex or triplex ddPCR assays. In order to quantify meat proportions, the percent ratio of *β-actin* copy number/*myostatin* copy number was calculated. The ratios of *β-actin*/*myostatin* were then plotted on the y-axis against the percent weight of meat matrices on the x-axis to show whether or not the fraction from the copy numbers of *β-actin*/*myostatin* gene was related to the percentage of mass fractions (Figure 5). The result showed that there were no statistically significant differences among the assays to quantify the proportion of pork DNA in the mixed meat (*P*> 0.05) (Figure 5A). It was found that the relationship between the percentage of gene copy ratio and the mass fraction was linear for all the three ddPCR assays for pork target (Figure 5A) and vice versa for beef mass in meat mixture when using the triplex assay (Figure 5B). The correlation coefficients (R^2^) for pork singleplex, duplex and triplex assays were 0.9988, 0.9996 and 0.9996, respectively whereas it was 0.9946 for the beef triplex assay. The coefficient of variation (CV) of the given result increased with a poor level of precision when the percentage of pork content decreased (Figure 6A). The same increasing trend of bias with a poor trueness was also observed with the decreasing percentage of pork content (Figure 6B). The CV and bias from singleplex ddPCR at 0.01% pork in beef were higher than those of the duplex and triplex assays. This might be due to the subsampling error associated with the low concentrations of the target molecules. This type of error was also presented when using qPCR (Köppel et al 2020; Soares et al 2010; Taylor et al. 2019). Simultaneous PCR reactions in duplex and triplex assays could possibly reduce technical errors, reagents, and time requirement (Köppel et al 2020; Whale et al. 2016). In addition, the performance of the myo assay has been shown not only to be suitable for quantifying the total amount of meat, but also for confirming the quality of the extracted nucleic acid for a reliable exclusion of false-negative detections (Laube et al. 2003; Nixon et al. 2015). The internal control plasmid DNA has been previously used for quality control by Shehata et al. (2017). However, this control has to be aware of the cross reactivity and the competition to target sequences. Moreover, adding the internal control plasmid may risk losing plasmid DNA during the extraction processes, resulting in not being representative as the real internal control (Shehata et al. 2017). The *myostatin* could be an alternative real internal control to ensure reliability, normalised variabilities and safeguard against false negatives. However, care should be taken when using the DNA based method in the quantification purpose as a big different genome size of meat species may affect the accuracy of ratio quantification; for example, a difference in the genome size between pork (2,800 Mb) and chicken (200-5 Mb) (Burt 2005; Groenen et al. 2012). Further study needs to be carried out to investigate the impact of the genome size in the ratio of DNA quantification.

**Figure 5.**
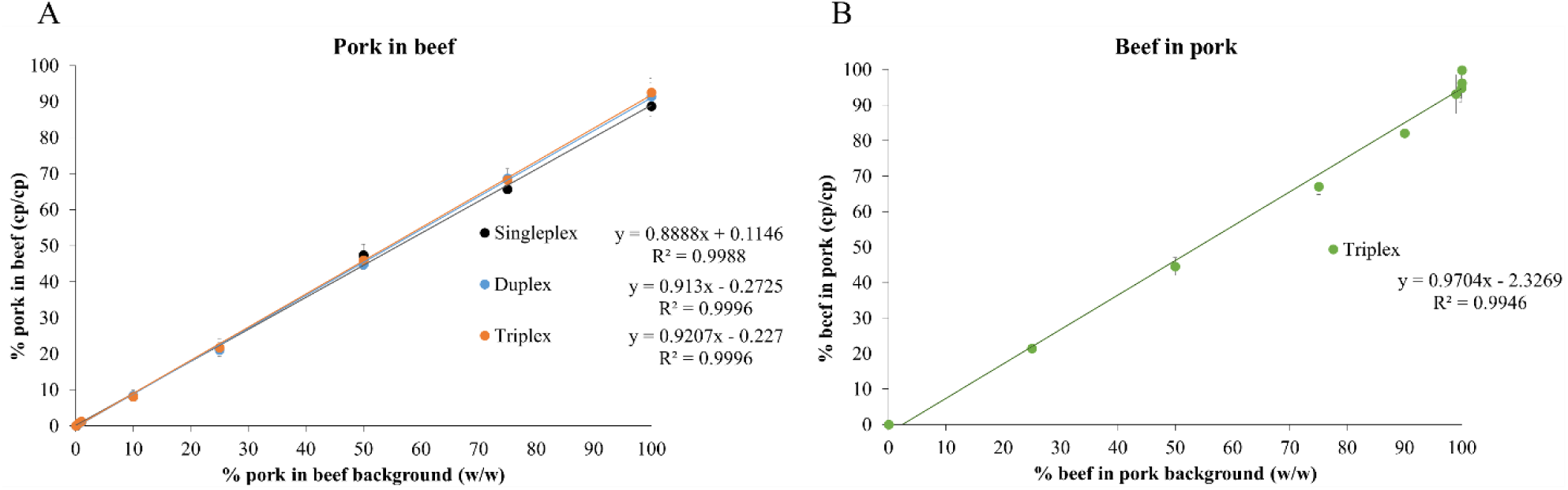
Linear regression between the percentage of ddPCR output ratio (cp/cp) and the percentage of expected pork adulteration with beef by gravimetric balance method (w/w). A: % pork in a beef background that was measured by singleplex (black), duplex (blue) and triplex (orange) ddPCR assays (y-axis) compared with assigned value (x-axis). B: For triplex ddPCR assay for beef quantification in pork background.

**Figure 6.**
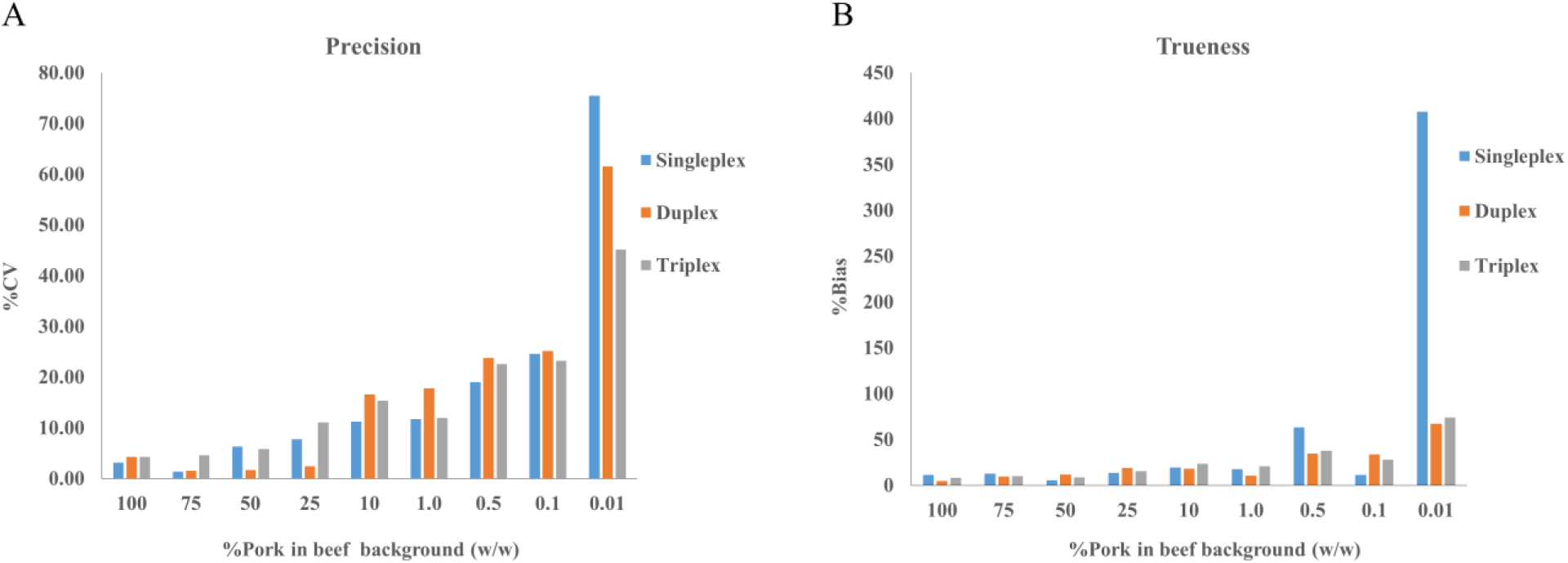
Accuracy of singleplex (blue), duplex (orange) and triplex (grey) ddPCR assays. A: The percentage of coefficient of variation (%CV) and B: the percent of bias (%bias) expressed precision and trueness, respectively.

In order to determine the LOD and LOQ, the DNA from 10 independent replicates for each ratio of pork in beef matrices (10%, 1%, 0.5%, 0.1%, 0.01% and 0% (w/w) pork in beef) were isolated. LOD was defined as the lowest percentage of pork content that could be reliably detected. Although the proportion of pork in beef was decreased to 0.01%, the developed ddPCR assays can still estimate pork containing in samples. With the limit of gravimetric balance, the LOD of all developed pork detection assays by ddPCR was ≤ 0.01% (Table 2). LOQ was defined as the lowest percentage of pork content that was precisely quantified with confidence (<25%CV) (Cai et al. 2017; Deprez et al. 2016; Košir et al. 2017). Therefore, 0.1% pork contamination in beef was the LOQ for singleplex, duplex and triplex ddPCR assays (Table 2).

**Table 2.**
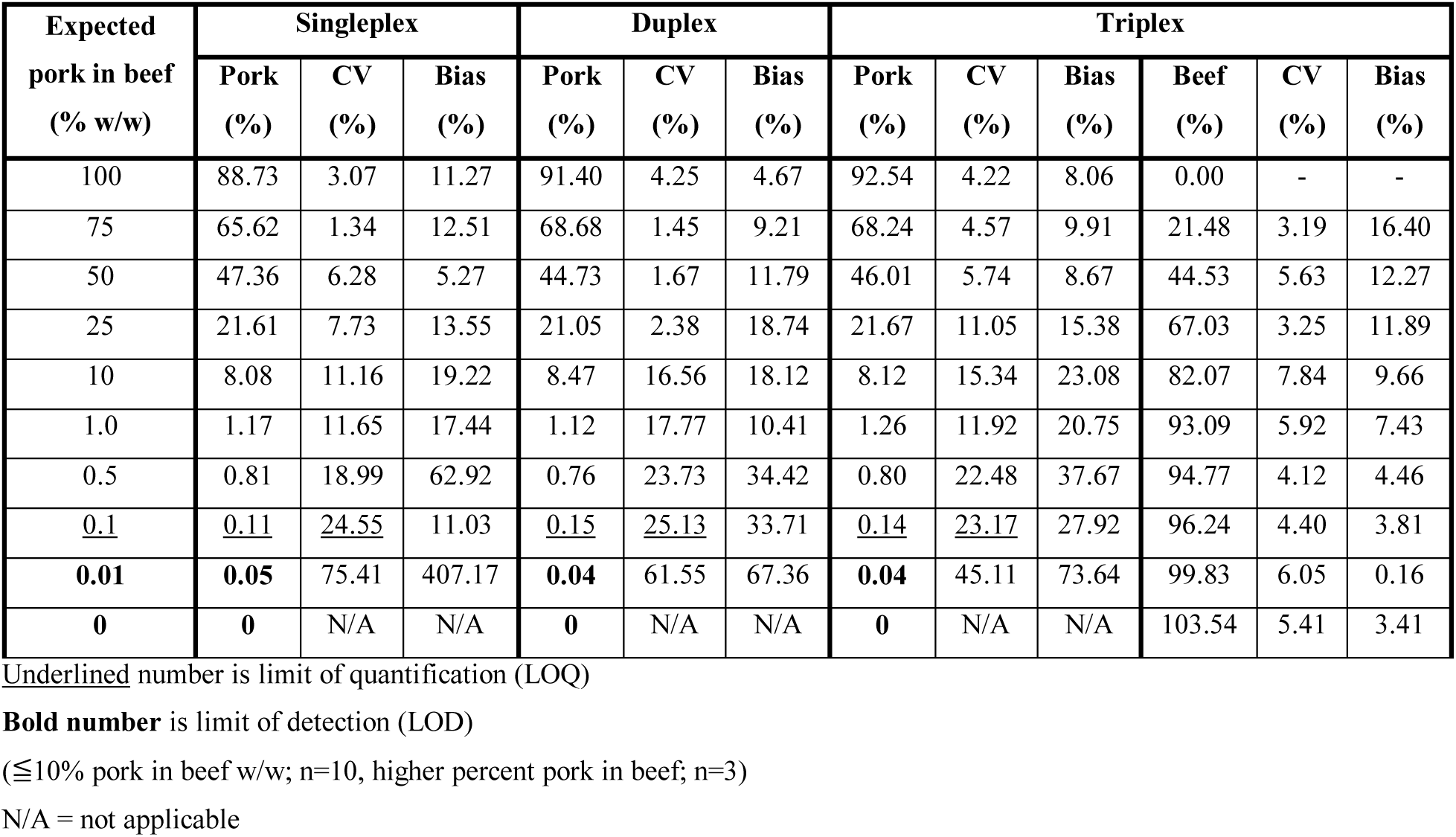
Accuracy (%CV and % bias represent precision and trueness, respectively) of the developed ddPCR assays in measuring pork and beef meat.

| Expected<br>pork in beef<br>(% w/w) | Singleplex |  |  | Duplex |  |  | Triplex |  |  |  |  |  |
| --- | --- | --- | --- | --- | --- | --- | --- | --- | --- | --- | --- | --- |
|  | Pork<br>(%) | CV<br>(%) | Bias<br>(%) | Pork<br>(%) | CV<br>(%) | Bias<br>(%) | Pork<br>(%) | CV<br>(%) | Bias<br>(%) | Beef<br>(%) | CV<br>(%) | Bias<br>(%) |
| 100 | 88.73 | 3.07 | 11.27 | 91.40 | 4.25 | 4.67 | 92.54 | 4.22 | 8.06 | 0.00 | - | - |
| 75 | 65.62 | 1.34 | 12.51 | 68.68 | 1.45 | 9.21 | 68.24 | 4.57 | 9.91 | 21.48 | 3.19 | 16.40 |
| 50 | 47.36 | 6.28 | 5.27 | 44.73 | 1.67 | 11.79 | 46.01 | 5.74 | 8.67 | 44.53 | 5.63 | 12.27 |
| 25 | 21.61 | 7.73 | 13.55 | 21.05 | 2.38 | 18.74 | 21.67 | 11.05 | 15.38 | 67.03 | 3.25 | 11.89 |
| 10 | 8.08 | 11.16 | 19.22 | 8.47 | 16.56 | 18.12 | 8.12 | 15.34 | 23.08 | 82.07 | 7.84 | 9.66 |
| 1.0 | 1.17 | 11.65 | 17.44 | 1.12 | 17.77 | 10.41 | 1.26 | 11.92 | 20.75 | 93.09 | 5.92 | 7.43 |
| 0.5 | 0.81 | 18.99 | 62.92 | 0.76 | 23.73 | 34.42 | 0.80 | 22.48 | 37.67 | 94.77 | 4.12 | 4.46 |
| <u>0.1</u> | <u>0.11</u> | <u>24.55</u> | 11.03 | <u>0.15</u> | <u>25.13</u> | 33.71 | <u>0.14</u> | <u>23.17</u> | 27.92 | 96.24 | 4.40 | 3.81 |
| **0.01** | **0.05** | 75.41 | 407.17 | **0.04** | 61.55 | 67.36 | **0.04** | 45.11 | 73.64 | 99.83 | 6.05 | 0.16 |
| **0** | **0** | N/A | N/A | **0** | N/A | N/A | **0** | N/A | N/A | 103.54 | 5.41 | 3.41 |

Underlined number is limit of quantification (LOQ)
**Bold number** is limit of detection (LOD)
( $\leq 10\%$ pork in beef w/w; n=10, higher percent pork in beef; n=3)
N/A = not applicable

### Analysis of processed foods

As the triplex ddPCR assay has the potential to reduce cost and time with high accuracy and reliability as shown this study, the triplex ddPCR assay was used to evaluate nine commercial processed food products and three autoclave treated meat samples (Table 3). The results demonstrated that six samples were identified as pork in the products and five products showed positive signals of beef, while only one sample (shrimp) did not detect the *myostatin* gene as expected (Table 3). Other works also found the qPCR methods have the capability for detecting traces of meat species in processed food under various processing conditions (Ali et al. 2012; Barakat et al. 2014; Che Man et al. 2012; Naaum et al. 2018; Yusop et al. 2012). For quantitative detection, eight of the nine samples showed a successful quantification with the percentage of pork or beef close to the declared percentage on their labels for commercial meat products, or to the true value of the weight proportions for mimicking processed foods (Table 3). Interestingly, there was one product, pork sausage A, showing the detection result of the percentage of pork content by only 30.40%± 1.78%, which was approximately 3-times lower than the declaration on its label (87%).

**Table 3.**
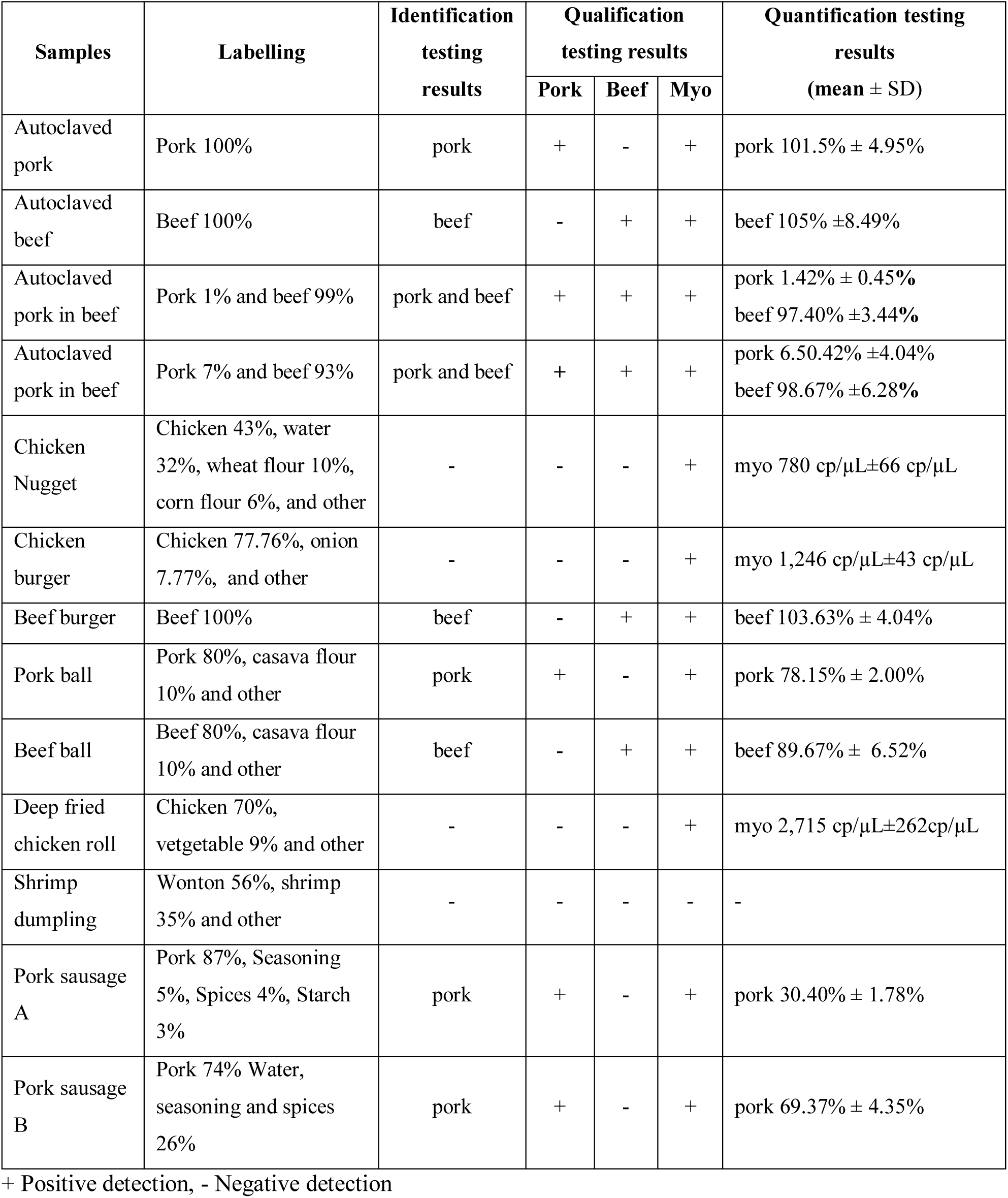
Determination of pork or beef meat percentage in highly processed foods (autoclaved meat) and commercial processed foods (products from Thailand) by triplex ddPCR assay. Three biological replicates and three technical replicates were presented.

When comparing the autoclaving to other cooking methods such as boil, grill or stream, the extracted DNA from the autoclave treated meat samples were clearly fragmented (approximately 80-500 bp) than other cooking methods (Figure S4). Although, the degraded DNA was observed, the DNA fragment size was still in a range of the target template with the same DNA degradation pattern among the meat through the samples (Figure S4). Consequently, these might not affect the quantification of the ratio in the highly processed foods by the ddPCR assays. Moreover, in the ddPCR system, the restriction enzyme is recommended for digesting the input genomic DNA as the viscosity of template can interfere the partition samples (Yukl et al. 2014). As a result, the DNA fragment from highly processed foods may appropriate for ddPCR systems without the treatment with restriction enzyme. This work provides the developed ddPCR assays that can be employed to accurately determine not only qualitative, but also quantitative data of simultaneously pork, beef and *myostatin* at a time in the processed meat samples without the requirement of the calibration curve. We, therefore, believe that the ddPCR assays developed in this study are suitable and could be very useful for investigating the mislabelling of meat content which could be either unwittingly or deliberately added.

## Conclusions

The results described in this work provide the evidence that the developed ddPCR assays showed their ability of qualitative and quantitative measurements of pork in meat mixtures based on DNA content with high sensitivity, specificity and accuracy, which the ratio of copy number DNA can be directly related to the mass fraction without the calibration curve or constant number. In the developed duplex and triplex ddPCR assays, the introduction of the cross species (*myostatin*) gene for quantifying the total amount of meat background was shown to minimise the effects from technical variations and bias. The myo assay also provided the required confidence in processed food analysis highlighting its suitability to being used as an internal and quality control. In addition, the triplex assay was able to quantify pork, beef and *myostatin* genes simultaneously in one PCR reaction. The triplex assay demonstrated good performance when applied with autoclaved meat and commercial food products, indicating that developed ddPCR assays have great potential to be utilised as a standard method in the determination of pork and/or beef fractions to aid regulations in controlling food adulteration.

