## Appendix A supplementary data for "Accurate determination of meat mass fractions using DNA measurements for quantifying meat adulteration"

***Figure S1.***

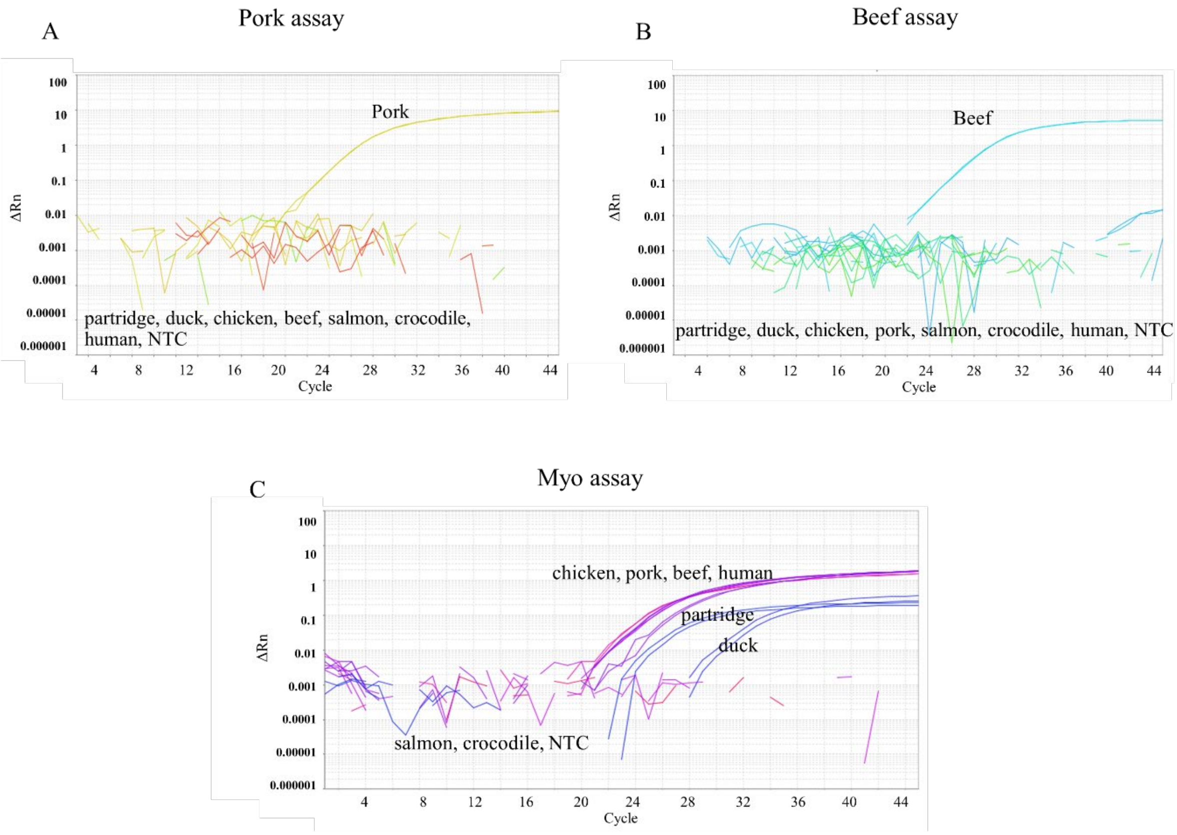


**Figure S1.** Amplification plots of qPCR for specificity test of primers and probes to DNA extracted from different animal species. Pork (A), beef (B) and myo (C) assays were challenged with DNAs extracted from chicken, pork, beef, duck, partridge, salmon, crocodile and human DNA. NTC is no-DNA-template control.

*Figure S2.*

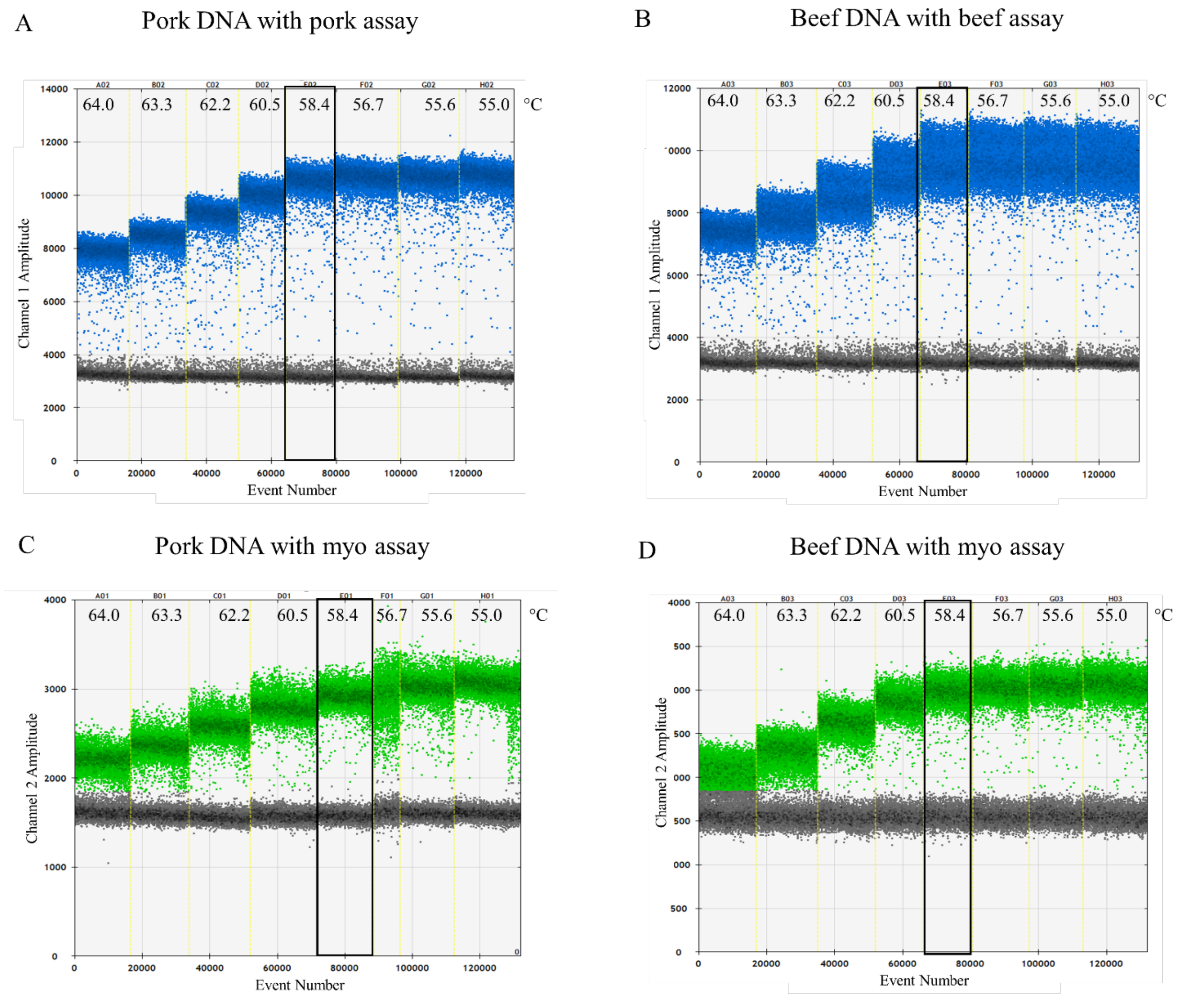


**Figure S2.** Optimisation of annealing temperature of primers and probes by ddPCR system. The PCR reactions were observed using a thermal gradient PCR ranging from 55 to 64 ºC. The figure shows the discrimination of droplet populations, black: negative droplets, blue: positive droplets with *β-actin* gene targets in pork (A) and beef (B) DNA by pork and beef assays, respectively. Green: positive droplets with *myostatin* gene target in pork (C) and beef (D) DNA by myo assay.

*Figure S3.*

| Assay/ Target/ Mean PCR EF | Amplification Plots | Calibration Curve |
|---|---|---|
| Pork/pork 94.93 % ±3.50% | A 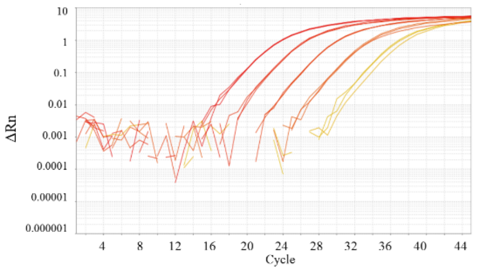  | E 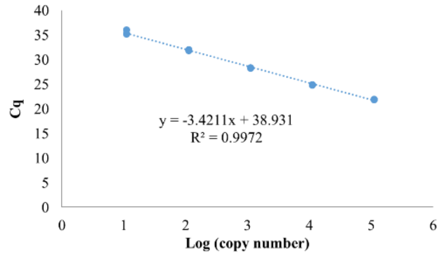  |
| Beef /beef 96.38 %±1.54% | B 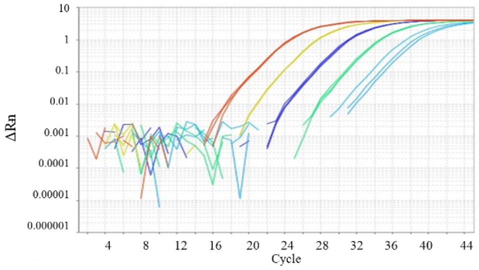  | F 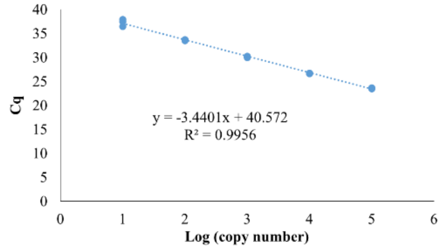  |
| Myo /pork 97.23 %±5.16% | C 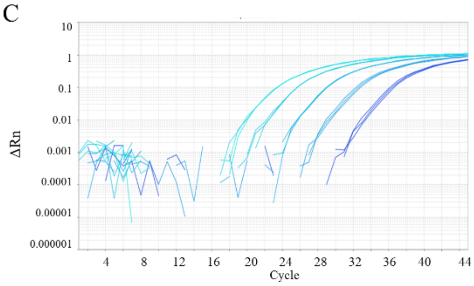  | G 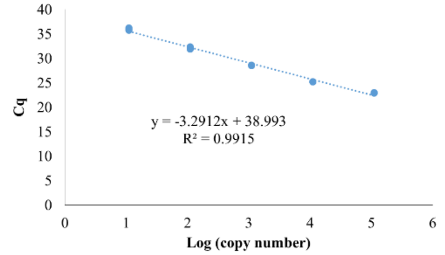  |
| Myo /beef 101.58 %±4.34% | D 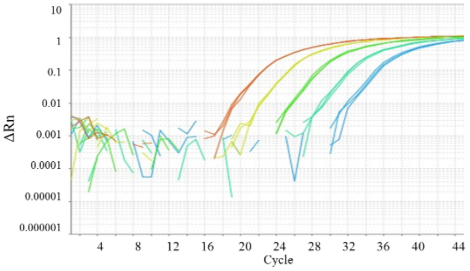  | H 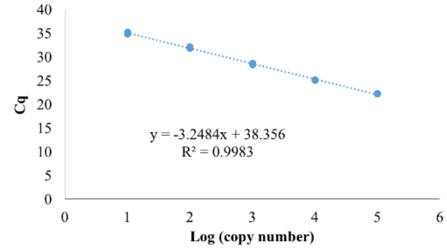  |

**Figure S3**. The amplification plots for each assay. The amplification plots of qPCR for 10-fold dilution series from $10^5$ to 10 copies/µL of pork DNA by the pork assay (A), beef DNA by the beef assay (B), pork

DNA by the myo assay (C) and beef DNA by the myo assay (D). The E-H graphs show the linear regression line derived from A-D graphs; pork DNA by the pork assay (E), beef DNA by the beef assay (F), pork DNA by the myo assay (G), and beef DNA by the myo assay (H). Three replicates were performed for each assay. PCR efficiency (PCR EF) can be calculated from Efficiency = ($10^{-1/slope}$-1), where the slope can be obtained from the linear regression equation. Pure pork and beef genomic DNAs were used while PCR efficiencies were derived from three independent experiments.

***Figure S4.***

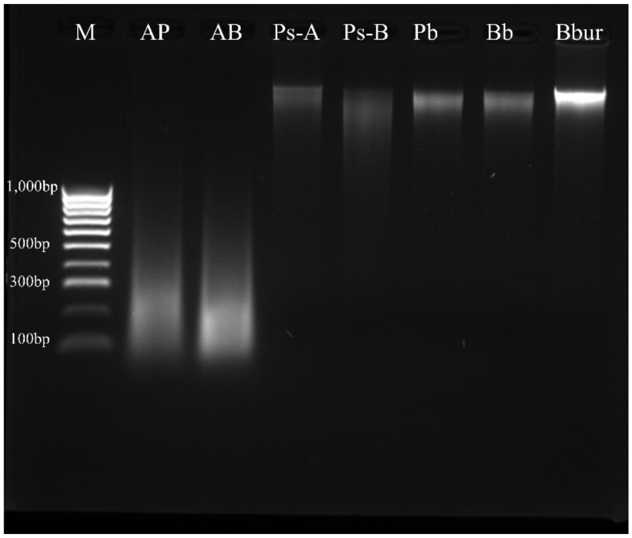


**Figure S4.** Example of 1.5 % agarose gel electrophoresis of extracted DNA derived from proceeded foods. M: marker, AP: autoclaved 100% pork, AB: autoclaved 100% beef, Ps-A: pork sausage A, Ps-B: pork sausage B, Pb: pork ball, Bb: beef ball, and Bbur: beef burger.

**Table S1. Digital MIQE checklist**

| ITEM TO CHECK | IMPORTANCE | CHECKLIST | COMMENTS/WHERE? |
|---|---|---|---|
| **EXPERIMENTAL DESIGN** | | | |
| Definition of experimental and control Groups | E | YES | See the part of materials and methods (known sample used as the control groups and commercial food products used as experimental groups) |
| Number within each group. | E | YES | Indicated in the heading of Table 2 and Table 3 |
| Assay carried out by core lab or Investigators' lab? | D | N/A | |
| Power analysis | D | N/A | |
| **SAMPLE** | | | |
| Description | E | YES | See the part of materials and methods |
| Volume or mass of sample processed | E | YES | The matrix of mixed meat using mass fraction (w/w). |
| Microdissection or macrodissection | E | N/A | |
| Processing procedure | E | N/A | |
| If frozen - how and how quickly? | E | N/A | |
| If fixed - with what, how quickly? | E | N/A | |
| Sample storage conditions and duration (especially for FFPE samples) | E | YES | -80 ºC for long storage and -20 ºC for short storage |
| **NUCLEIC ACID EXTRACTION** | | | |
| Quantification - instrument/method | E | YES | Experimental Section |
| Storage conditions: temperature, concentration, duration, buffer | E | YES | -80 ºC for long storage (for up to 1 year) and -20 ºC for short storage (4 week) in TE buffer |
| DNA or RNA quantification | E | YES | DNA |
| Quality/integrity- instrument/method; e.g. RIN/RQI and trace or 3':5' | E | YES | Using Gel electrophoresis, and Nanodrop (Available on request) |
| Template structural information | E | N/A | |

| | | | |
|---|---|---|---|
| Template modification (digestion, sonication, pre-amplification etc.) | E | N/A | |
| Template treatment (initial heating or chemical denaturation) | E | N/A | |
| Inhibition dilution or spike | E | YES | Each assay was performed on dilutions of the template. All assays displayed a linear relationship between amplification and copy number |
| DNA contamination assessment of RNA sample | E | N/A | |
| Details of DNase treatment where performed | E | N/A | |
| Manufacturer of reagents used and catalogue number | D | YES | DNeasy Mericon Food Kit (Qiagen, Hilden, Germany) |
| Storage of nucleic acid: temperature, concentration, duration, buffer | E | YES | Samples were stored in TE Buffer at -80ºC for up to 1 year. |
| **REVERSE TRANSCRIPTION (If necessary)** | | | |
| cDNA priming method + concentration | E | N/A | |
| One or two step protocol | E | N/A | |
| Amount of RNA used per reaction | E | N/A | |
| Detailed reaction components and conditions | E | N/A | |
| RT efficiency | D | N/A | |
| Estimated copies measured with and without addition of RT* | D | N/A | |
| Manufacturer of reagents used and catalogue number | D | N/A | |
| Reaction volume (for two step reverse transcription reaction) | D | N/A | |
| Storage of cDNA: temperature, concentration, duration, buffer | D | N/A | |
| **dPCR TARGET INFORMATION** | | | |
| Sequence accession number | E | YES | See Table 1 |
| Location of amplicon | D | YES | Available on request |
| Amplicon length | E | YES | See Table 1 |
| In silico specificity screen (BLAST, etc) | E | YES | Previous studies Köppel et al. 2011, Laube et al. 2003 |

| | | | |
|---|---|---|---|
| Pseudogenes, retropseudogenes or other homologs? | D | N/A | |
| Sequence alignment | D | YES | Available on request |
| Secondary structure analysis of amplicon and GC content | D | YES | The assays were previously used from publications (Table1). |
| Location of each primer by exon or intron (if applicable) | E | N/A | |
| Where appropriate, which splice variants are targeted? | E | N/A | |
| **dPCR OLIGONUCLEOTIDES** | | | |
| Primer sequences and/or amplicon context sequence** | E | YES | See Table 1 |
| RTPrimerDB Identification Number | D | N/A | |
| Probe sequences** | D | YES | See Table 1 |
| Location and identity of any modifications | E | YES | See Table 1 |
| Manufacturer of oligonucleotides | D | YES | Primers and dual labelled probes were manufactured by Macrogen, Korea. |
| Purification method | D | YES | Desalted for primers and HPLC for probes |
| **dPCR PROTOCOL** | | | |
| Complete reaction conditions | E | YES | Experimental Section |
| Reaction volume and amount of RNA/cDNA/DNA | E | YES | Experimental Section |
| Primer, (probe), Mg++ and dNTP concentrations | E | YES | Experimental Section; Manufacturer's proprietary |
| Polymerase identity and concentration | E | YES | Experimental Section; concentration is Manufacturer's proprietary |
| Buffer/kit Catalogue No and manufacturer | E | YES | ddPCR Supermix for probes (no dUTP), Cat#186-3024, BioRad |
| Exact chemical constitution of the buffer | D | NO | Manufacturers' proprietary |
| Additives (SYBR Green I, DMSO, etc.) | E | N/A | |
| Plates/tubes Catalogue No and manufacturer | D | N/A | |
| Complete thermocycling parameters | E | YES | Experimental Section |
| Reaction setup | D | YES | Experimental Section |

| | | | |
|---|---|---|---|
| Gravimetric or volumetric dilutions (manual/robotic) | D | YES | Volumetric dilutions (manual) |
| Total PCR reaction volume prepared | D | YES | 25 µL |
| Partition number | E | YES | Average 15774 |
| Individual partition volume | E | YES | 0.85 µL |
| Total volume of the partitions measured (effective reaction size) | E | YES | 20 µL |
| Partition volume variance/standard deviation | D | N/A | |
| Comprehensive details and appropriate use of controls | E | YES | Known %pork in beef (w/w) background were used |
| Manufacturer of dPCR instrument | E | YES | QX200 Bio-Rad |
| **dPCR VALIDATION** | | | |
| Optimisation data for the assay | D | YES | Available on request |
| Specificity (when measuring rare mutations, pathogen sequences etc.) | E | YES | This was checked using qPCR (Figure S1) |
| Limit of detection of calibration control | D | YES | See Table 2 |
| If multiplexing, comparison with singleplex assays | E | YES | See Table 2 |
| **DATA ANALYSIS** | | | |
| Average copies per partition (λ or equivalent ) | E | YES | (vary depend on concentrations) Available on request |
| dPCR analysis program (source, version) | E | YES | QuantaSoft v.1.7.4.0917 (Bio-rad) |
| Outlier identification and disposition | E | N/A | Used Grubbs's test (Available on request) |
| Results of NTCs | E | YES | All NTCs gave negative results and also see Figure 1 |
| Examples of positive(s) and negative experimental results as supplemental data | E | YES | See Figure 1 |
| Where appropriate, justification of number and choice of reference genes | E | N/A | |
| Where appropriate, description of normalisation method | E | YES | Experimental Section |
| Number and concordance of biological replicates | D | N/A | |
| Number and stage (RT or qPCR) of technical replicates | E | YES | Three replicates |

| | | | |
|---|---|---|---|
| Repeatability (intra-assay variation) | E | YES | See Table 2 |
| Reproducibility (inter-assay/user/lab etc. variation ) | D | N/A | |
| Experimental variance or confidence interval*** | E | YES | See Table 2 |
| Statistical methods used for analysis | E | YES | *Two-way ANOVA* |
| Data submission using RDML | D | N/A | |

dMIQE checklist for authors, reviewers and editors. All essential information (E) must be submitted with the manuscript. Desirable information (D) should be submitted if possible.

* Assessing the absence of DNA using a no RT assay (or where RT has been inactivated) is essential when first extracting RNA. Once the sample has been validated as DNA-free, inclusion of a no-RT control is desirable, but no longer essential.

** Disclosure of the primer and probe sequence is highly desirable and strongly encouraged. However, since not all commercial pre-designed assay vendors provide this information when it is not available assay context sequences must be submitted (48)

*** When single dPCR experiments are performed, the variation due to counting error alone should be calculated from the binomial (or suitable equivalent) distribution.
